# Nowcasting by Bayesian Smoothing: A flexible, generalizable model for real-time epidemic tracking

**DOI:** 10.1101/663823

**Authors:** Sarah F. McGough, Michael A. Johansson, Marc Lipsitch, Nicolas A. Menzies

## Abstract

Delays in case reporting are common to disease surveillance systems, making it difficult to track diseases in real-time. “Nowcast” approaches attempt to estimate the complete case counts for a given reporting date, using a time series of case reports that is known to be incomplete due to reporting delays. Modeling the reporting delay distribution is a common feature of nowcast approaches. However, many nowcast approaches ignore a crucial feature of infectious disease transmission—that future cases are intrinsically linked to past reported cases—and are optimized to a single application, which may limit generalizability. Here, we present a Bayesian approach, NobBS (Nowcasting by Bayesian Smoothing) capable of producing smooth and accurate nowcasts in multiple disease settings. We test NobBS on dengue in Puerto Rico and influenza-like illness (ILI) in the United States to examine performance and robustness across settings exhibiting a range of common reporting delay characteristics (from stable to time-varying), and compare this approach with a published nowcasting package. We show that introducing a temporal relationship between cases considerably improves performance when the reporting delay distribution is time-varying, and we identify trade-offs in the role of moving windows to accurately capture changes in the delay. We present software implementing this new approach (R package “NobBS”) for widespread application.

**Significance:** Achieving accurate, real-time estimates of disease activity is challenged by delays in case reporting. However, approaches that seek to estimate cases in spite of reporting delays often do not consider the temporal relationship between cases during an outbreak, nor do they identify characteristics of robust approaches that generalize to a wide range of surveillance contexts with very different reporting delays. Here, we present a smooth Bayesian nowcasting approach that produces accurate estimates that capture the time evolution of the epidemic curve and outperform a previous approach in the literature. We assess the performance for two diseases to identify important features of the reporting delay distribution that contribute to the model’s performance and robustness across surveillance settings.

## Introduction

Effective public health action relies on surveillance that is timely and accurate, especially in disease outbreaks(1, 2). Specifically, surveillance provides the information required to assess risks, prioritize and allocate resources to public health threats, deploy and discontinue interventions to interrupt transmission, and monitor the impact of those interventions. Ideally, disease surveillance systems should closely track the often fast-changing circumstances of outbreaks, distinguishing true changes in the dynamics from artifacts of reporting.

Despite the importance of timely surveillance data, substantial challenges exist to collect and report case information in real time. Multiple features of the disease and surveillance system contribute to reporting delays, including: delays in symptoms onset after infection; delays in medical care-seeking after onset; delays in providers obtaining and reporting diagnostic information; level of awareness of disease activity influencing care-seeking and reporting; and system-level processing delays, a result of complex and multi-tiered disease reporting and communication systems interacting at multiple administrative levels(3). Reporting delays can be further exacerbated in resource-constrained settings. As a consequence, surveillance data are typically not complete until weeks or months after infections have actually occurred, providing an incomplete picture of current disease activity.

Nowcasting, or “predicting the present,” is an approach to mitigate the impact of reporting delays. With origins in the insurance claims and actuarial literature(4, 5), nowcast models aim to estimate the number of occurred-but-not-yet-reported events (e.g. insurance claims, disease cases) at any given time based on an incomplete set of reports. In public health settings, nowcasting approaches have been explored for AIDS in the 1980s and 1990s(6–8) as a consequence of the long incubation period from HIV infection until development of AIDS. More recently, nowcasting has been applied to infectious disease outbreaks such as foodborne illness outbreaks(9, 10). These studies draw principally on survival analysis and actuarial techniques to model the reporting delay and draw inferences based on historical patterns. A majority of studies have strictly focused on modeling the reporting delay distribution—a legacy of the actuarial techniques giving rise to many of these approaches—and generally neglect a key feature of outbreaks: that future cases are intrinsically linked to past reported cases, a fact that creates potentially strong autocorrelation in the true number of cases over short time intervals. In other words, the infectious disease transmission process provides an additional signal of the number of cases to be expected in the near future that has not been included in common methods such as the reverse-time hazard model (11, 12) and the chain ladder method (13). However, proposals to extend the latter approach to state-space models that account for temporal relationships in reporting have existed in the literature since the development of these techniques(13–15) and have been applied in at least one infectious disease context(16). These developments are promising for disease surveillance, but it is critical to demonstrate performance in a diversity of settings as infectious disease nowcast models to date have largely focused on specific applications, not the common challenges that exist across many different diseases. In this investigation, we find that nowcasting is especially challenging when the proportion of cases reported the week they occur (delay 0) is low and reporting delays are highly variable; we know of no investigations that specifically identify models that perform well in these commonly occurring circumstances. As a result, the characteristics of robust and broadly-applicable models are difficult to identify. Additionally, several previous models have largely focused on providing point estimates of the number of cases. Point estimates may be helpful, but quantifying the uncertainty in those estimates is even more important in the context of infectious disease outbreaks because uncertainty is intrinsic, and accounting for plausible outcomes apart from the point prediction is critical.

Here, we introduce Nowcasting by Bayesian Smoothing (NobBS), a simple and flexible generalized Bayesian model for nowcasting infectious diseases in different settings. We demonstrate the robustness of this approach in two very different disease surveillance contexts and identify the conditions that favor its application, especially when the reporting delay distribution is time-varying. Specifically, NobBS allows for both uncertainty in the delay distribution and the time evolution of the epidemic curve, producing smooth, time-correlated estimates of cases. We demonstrate that NobBS performs well for weekly nowcasts of (1) dengue cases in Puerto Rico and (2) influenza-like illness (ILI) cases in the United States, requiring no disease-specific parameterization despite the two pathogens being very different (vector-borne vs. directly transmitted) and exhibiting substantially different reporting delays. Lastly, we test NobBS against a previous Bayesian nowcast method(9) and find that NobBS outperforms this benchmark for both diseases and multiple time periods. In particular, we show that while point estimates of the models are similar when time-to-report distributions are relatively fixed over time, NobBS improves the estimation of uncertainty and accommodates temporal variation in delay probabilities. We present an R package, “NobBS,” as a tool to complement both routine public health surveillance as well as forecasting efforts.

## Results

We developed a Bayesian approach to nowcast total case numbers using incomplete, time-stamped reported case data based on an estimated delay distribution, intrinsic autocorrelation from the transmission process, and historical case data. Generally, the approach learns from historical information on cases reported at multiple delays (e.g. no delay, 1-week delay, 2-week delay, etc.) from the week of case onset to estimate the reporting delay probability at each delay and the relationship between case counts from week-to-week, and uses this relationship to predict the number of not-yet-reported cases in the present. We tested this approach, NobBS, using two different infectious disease surveillance data sources: dengue surveillance in Puerto Rico, and national notifications of influenza-like illness (ILI) in the United States. Using all of the available data on case reporting delays up to the point of prediction, weekly dengue nowcasts were estimated for the time period December 23, 1991 through November 29, 2010 (989 weeks), and weekly ILI nowcasts were produced over the period June 30, 2014 through September 25, 2017 (170 weeks). For comparison, we generated weekly nowcasts over the same periods using an existing Bayesian approach, here referred to as the benchmark approach (9). To leverage a large amount of historical data to fit the nowcast model while also having a large window over which to assess nowcasts, we used a 104-week (approximately 2-y) moving window dengue and a 27-week (approximately 6-mo) moving window for ILI. Our primary outcome metric to assess nowcast performance was the logarithmic score, a proper score that evaluates the probability assigned to the observed outcome rather than error associated with a point prediction. For purposes of discussion, we reported the exponentiated form of the mean logarithmic score (the geometric mean of the assigned probabilities) to provide a metric on the scale of 0 (no certainty of the outcome) to 1 (complete certainty of the outcome). In addition, we estimated other metrics describing the performance of point estimates (mean absolute error (MAE), root mean square error (RMSE), and relative root mean square error (rRMSE)) and the 95% prediction interval (PI) coverage, and of these, focus on comparing the rRMSE and 95% PI coverage across approaches.

### Performance in forecasting weekly dengue and influenza incidence

Figs. 1-2 show weekly dengue and ILI nowcasts for NobBS and the benchmark approach over multiple seasons for both diseases. Table 1 summarizes the point and probability-based accuracy metrics for each, where higher accuracy is indicated by higher average scores, lower MAE, RMSE, and rRMSE, and lower distance from 0.95 for the 95% PI coverage. Because the NobBS model accounts for both under-reporting and the autocorrelated progression of transmission across successive weeks, it makes predictions even in weeks when there are no cases reported for the week. Conversely, the benchmark model does not make nowcasts for weeks in which there are no initial case reports (common in the dengue Puerto Rico data), hence the nowcasts in Figs. 1C and 2C appear as discontinuous lines. To compare models despite these differences, we report accuracy metrics between NobBS and the benchmark approach for both (1) the full time series of the data and (2) weeks when at least one case was reported in the first week, i.e. the subset of weeks for which both models could make predictions (Table 1). We also computed error metrics for the benchmark model for the full time series by assigning point estimates of 0 cases for nowcasts in weeks without predictions.

**Table 1.**
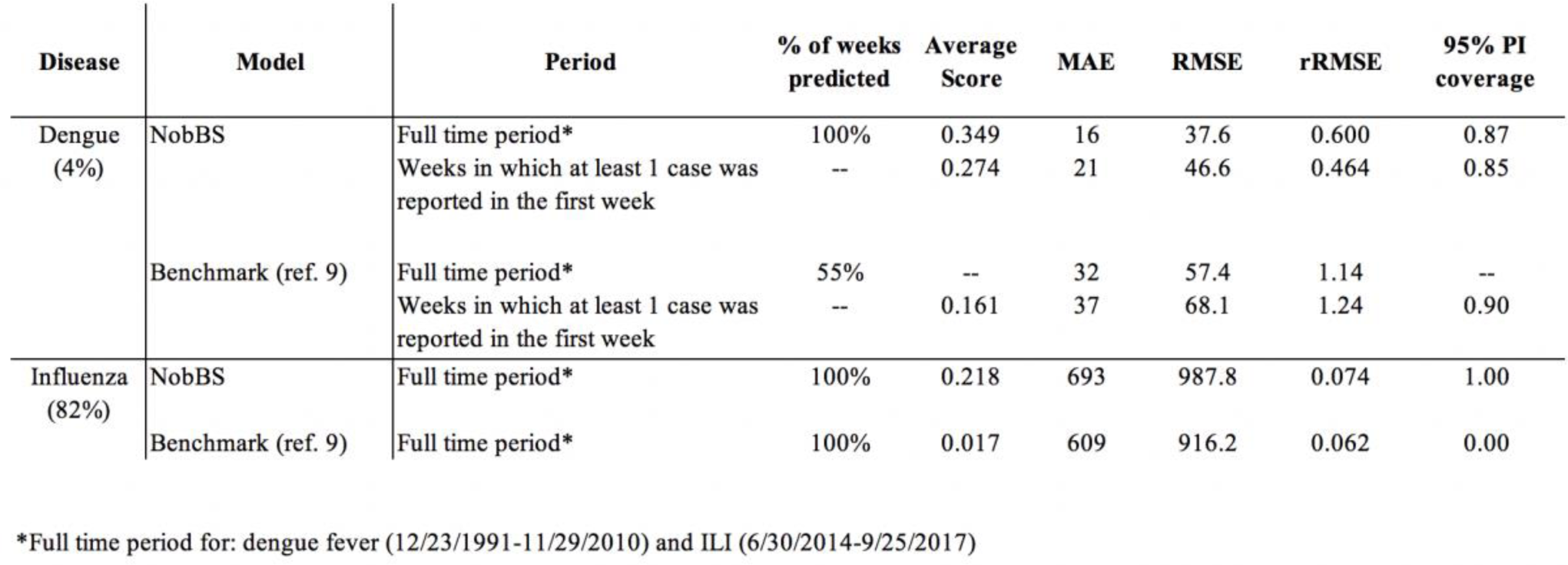
Performance measures for each nowcast approach and disease.

**Fig. 1.**
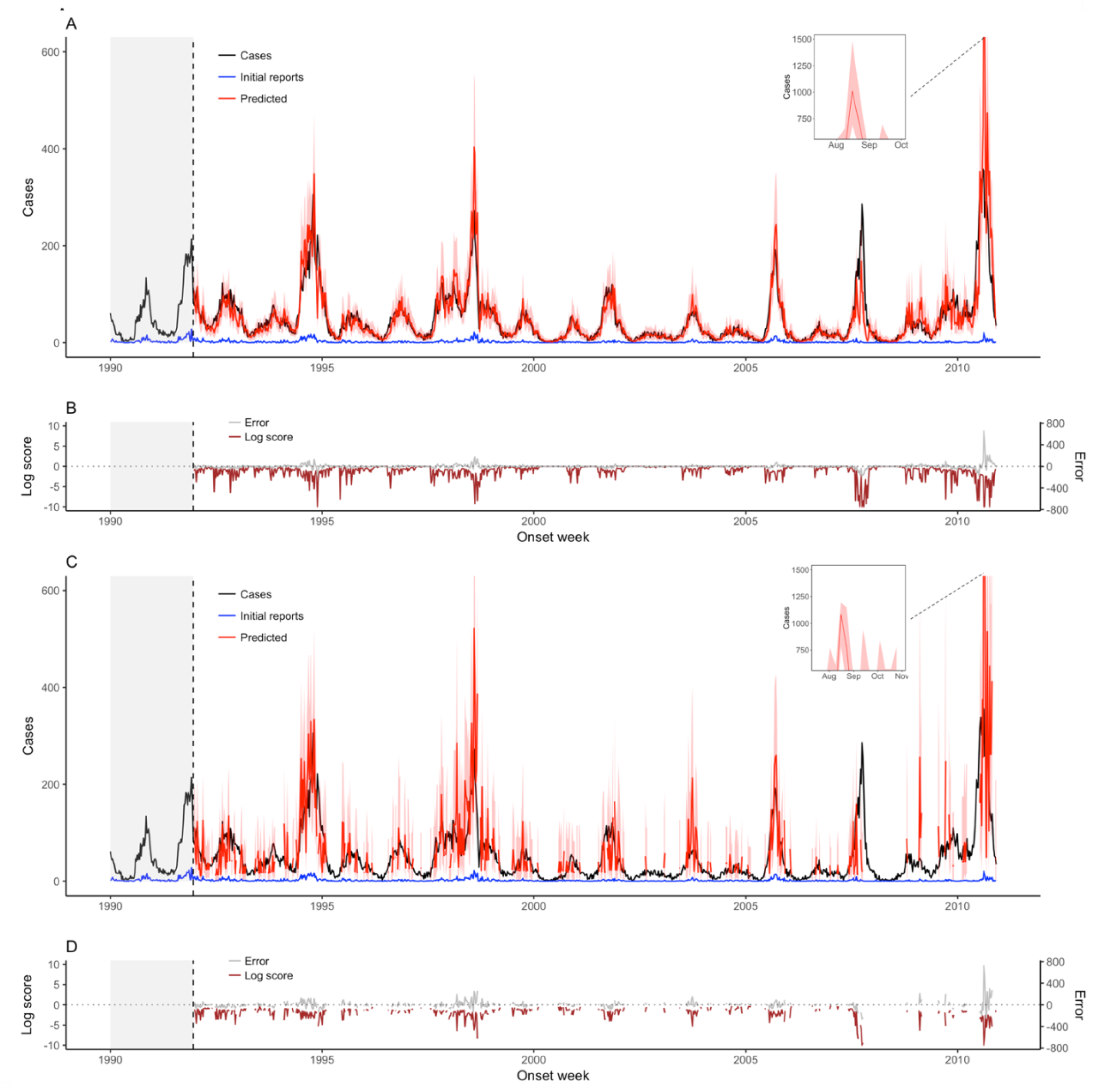
Weekly dengue fever nowcasts for December 23, 1991 through December 25, 2000 using a 2-year moving window. (A) NobBS nowcasts along with (B) point estimate and uncertainty accuracy, as measured by the log score and the prediction error, are compared to (C) nowcasts by the benchmark approach with (D) corresponding log scores and prediction errors. For nowcasting, the number of newly-reported cases each week (blue line) are the only data available in real-time for that week, and help inform the estimate of the total number of cases that will be eventually reported (red line), shown with 95% prediction intervals (pink bands). The true number of cases eventually reported (black line) is known only in hindsight and is the nowcast target. Historical information on reporting is available within a 104-week moving window (grey shade) and used to make nowcasts. The log score (brown line) and the difference between the true and mean estimated number of cases (grey line) are shown as a function of time.

**Fig. 2.**
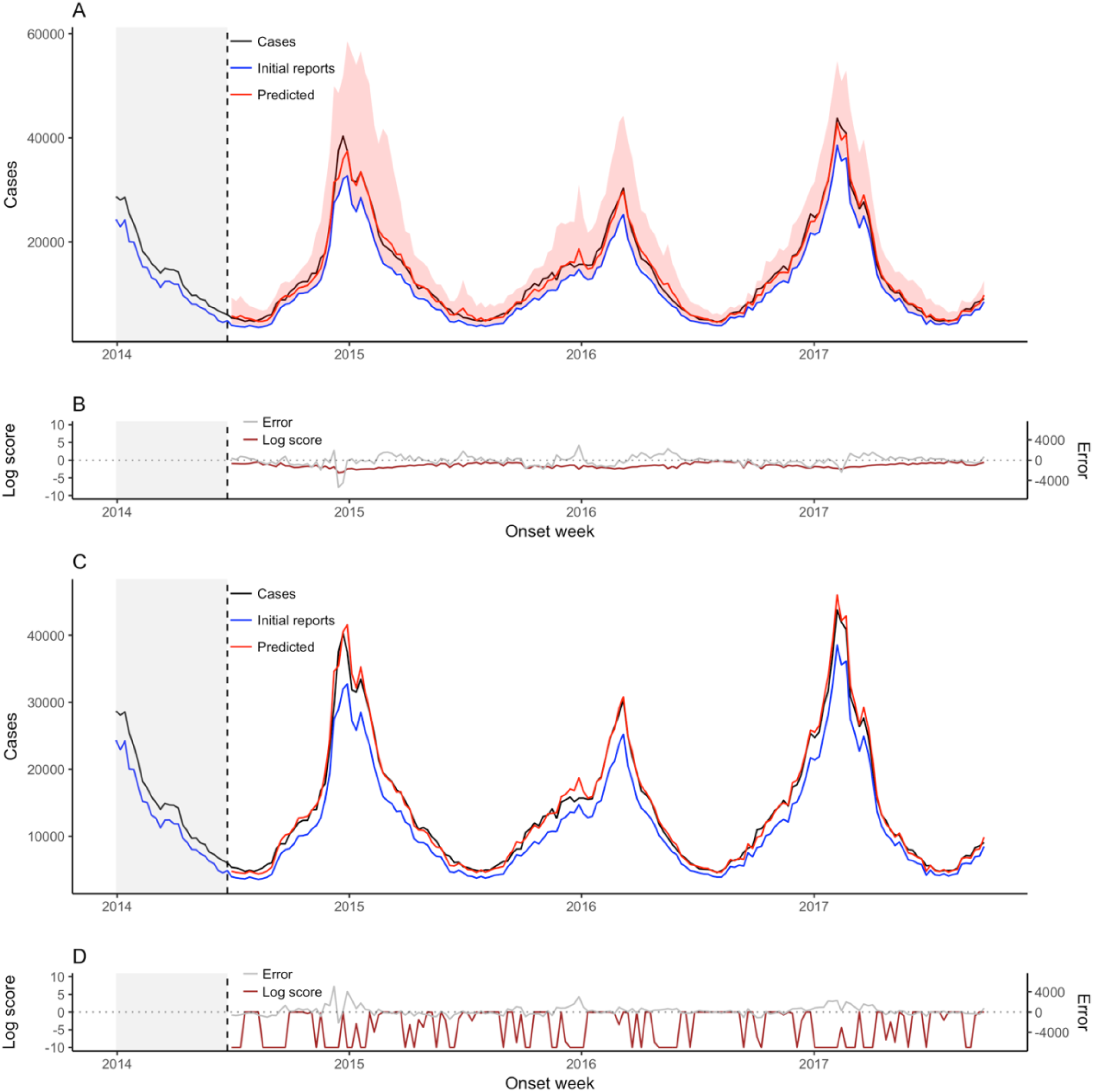
Weekly ILI nowcasts for June 30, 2014 through September 25, 2017 using a 6-month moving window. (A) NobBS nowcasts along with (B) point estimate and uncertainty accuracy, as measured by the log score and the prediction error, are compared to (C) nowcasts by the benchmark approach with (D) corresponding log scores and prediction errors. For nowcasting, the number of newly-reported cases each week (blue line) are the only data available in real-time for that week, and help inform the estimate of the total number of cases that will be eventually reported (red line), shown with 95% prediction intervals (pink bands). For the benchmark approach, the 95% prediction intervals are very narrow and are thus difficult to see. The true number of cases eventually reported (black line) is known only in hindsight and is the nowcast target. Historical information on reporting is available within a 27-week moving window (grey shade) and used to make nowcasts. The log score (brown line) and the difference between the true and mean estimated number of cases (grey line) are shown as a function of time.

The benchmark approach made predictions in only 55% of weeks in the dengue time series (Table 1). In this subset of weeks, the NobBS approach achieved relatively smooth and accurate tracking of the dengue time series (rRMSE = 0.464, average score = 0.274) despite low proportions of cases reported on the week of onset (Fig. 1A-B). The 95% PI coverage was 0.85, indicating that the 95% PI included the true number of cases for 85% of the nowcasts. In comparison, the benchmark approach produced substantially less accurate point estimates and slightly broader uncertainty intervals (rRMSE = 1.24, average score = 0.161, 95% PI coverage = 0.90) with greater fluctuation in nowcasts from week-to-week (Fig. 1C-D). Because many weeks in the dengue data were low incidence, assigning a prediction of 0 to the benchmark approach’s missing nowcasts improved its rRMSE to 1.14 in the full time series compared to 1.24 for the subset over which nowcasts were generated from the model, though NobBS still surpassed the benchmark model’s accuracy on this and all other metrics (Table 1).

Nowcast point estimates tracked the ILI time series well for both approaches, though point estimates had greater error by all measures for the NobBS approach (NobBS rRMSE = 0.074 vs. benchmark rRMSE = 0.062; Table 1). However, the NobBS approach produced considerably wider prediction intervals (Figs. 1C, 2C) resulting in both higher log scores (NobBS average score = 0.218 vs. benchmark average score = 0.017) and 100% coverage by the 95% prediction intervals compared to 0% coverage for the benchmark (Table 1).

To assess the degree of autocorrelation and related smoothness in the NobBS predictions, we calculated the 1-week lagged autocorrelation of predictions (ρ_a_) and compared this to the 1-week lagged autocorrelation of cases (ρ_c_). In addition, we computed metrics reflecting the accuracy of the approaches in capturing the *change* in cases from week-to-week: the mean absolute error of the change (MAEΔ) and the RMSE of the change (RMSEΔ) (Table 2). The magnitude of change was much larger for the ILI data than dengue data, with average absolute value change of 1,312.6 cases/week versus 9.8 cases/week, yet both showed high autocorrelation (ρ_c_ = 0.958 for dengue and ρ_c_ = 0.972 for ILI). Comparing the full time series, the nowcasts produced by NobBS exhibited high autocorrelation for both diseases (ρ_a_ = 0.876 for dengue, 0.973 for ILI) while the benchmark approach yielded lower autocorrelation for dengue nowcasts, comparatively (ρ_a_ = 0.631 for dengue, 0.970 for ILI). For dengue, over the weeks in which at least 1 case was initially reported, the NobBS approach achieved both lower mean absolute difference between predicted and observed changes in cases (NobBS MAEΔ = 23 vs. benchmark MAEΔ = 50) and lower RMSE of the change (NobBS RMSEΔ = 35.8 vs. benchmark RMSEΔ = 64.6). In addition, NobBS outperformed the benchmark approach over the full time series of dengue cases (Table 2). For ILI, however, the metrics for the weekly change were similar for the two approaches (Table 2).

**Table 2.**
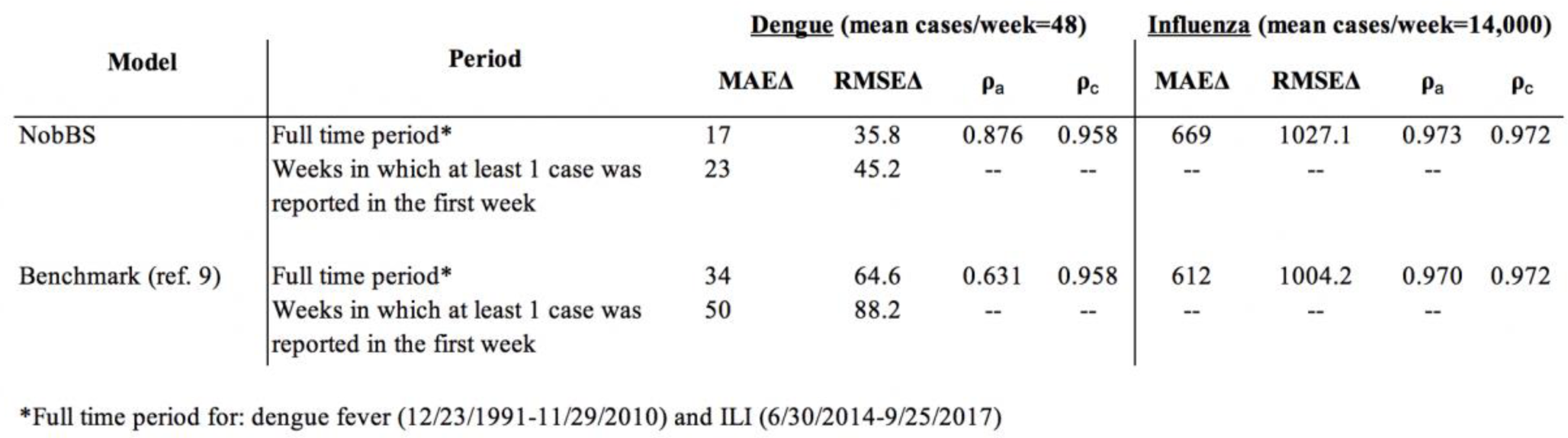

### Reporting delays impact nowcast performance

The delay distributions between the reporting systems are strikingly different (Figs. 1, 2, S1). In the case of the dengue surveillance system, which includes specimen collection and laboratory testing, only approximately 4% of cases were processed during the week of onset, on average. In contrast, the U.S. Outpatient Influenza-like Illness (ILI) Surveillance Network (ILINet) captures only syndromic data reported electronically, with over 80% of ILI cases reported, on average, the same week they present (i.e. with no delay). Overall, we observed that the accuracy of nowcast point estimates (rRMSE) was higher for the ILI data compared to dengue, which may be related to the high proportion of cases reported with 0-weeks delay in these data. Large weekly absolute changes in the number of initial case reports also appeared to be related to increased error, particularly for dengue, which had high fluctuations in the number of initial reports over time (Table S1, Fig. S2). Note that because of the difference in predictive distribution bin widths based on the number of cases that accrue for influenza vs. dengue (*Materials & Methods*), average scores are not comparable across diseases.

### NobBS improves nowcasting with varying reporting delays

Dengue and ILI also exhibit differences in the *trends* of reporting delay probabilities *over time*. For dengue, we observe a noisier, more time-varying probability of reporting for cases, with more extreme fluctuations in the proportion of initial reports compared to ILI cases, which show more constant (tighter ranges of) reporting probabilities from week-to-week (Fig. S3). Independent of the initial proportion of cases reported (high vs. low), we hypothesized that these trends (relatively constant vs. time-varying) are particularly impactful on the performance of the nowcast, and that relatively constant reporting probabilities, as seen in the ILI data, may be linked to the higher accuracy of these predictions.

To test the robustness of the model, we simulated ILI data using the final counts from the true dataset, but imposing a time-varying delay distribution; specifically, with faster initial reporting during weeks of high incidence (described in *Materials and Methods*). Using these simulated data, we found that NobBS was relatively robust to changes in reporting delays (Fig. S4, Table S2). In the context of stable reporting delays (original ILI data), NobBS performed comparably to the benchmark model (Fig. 2, Table 1). However, in the presence of simulated time-varying reporting delays, NobBS outperformed the benchmark in terms of confidence (NobBS average score = 0.06 vs. benchmark average score ≈ 0), point estimates (NobBS rRMSE = 0.302 vs. benchmark rRMSE = 0.621), and accuracy of the predicted change (Table S3). Such variations are a reality in many epidemics(17).

### Performance by year

The performance of ILI nowcasts across accuracy measures was relatively consistent by year, but there were fluctuations in the year-to-year performance of both approaches applied to dengue data (Table 3). Average scores tended to be high in years that experienced a very low number of dengue cases (e.g. 2000, 2002, 2004, 2006). The model was particularly effective at identifying periods of low incidence, with high probabilities assigned to the lowest outcome bin (0-25 cases, details in *Materials and Methods*) when the number of cases eventually reported was low (Fig. S5). On the other hand, during periods of high dengue activity, lower probabilities were assigned to the correct bin, reflecting greater uncertainty. Overall, NobBS outperformed the benchmark approach on all performance measures for each year (Table 3).

**Table 3.** Annual performance measures for each nowcast model, by disease. All predicted weeks for each model are compared.

| Disease | Year | Cases | NobBS |  |  |  | Benchmark (ref. 9) |  |  |  |
| --- | --- | --- | --- | --- | --- | --- | --- | --- | --- | --- |
|  |  |  | MAE | rRMSE | RMSE | Average Score | MAE | rRMSE | RMSE | Average Score |
| Dengue | 1992 | 3,570 | 15 | 0.271 | 19.7 | 0.262 | 27 | 0.473 | 33.2 | 0.154 |
|  | 1993 | 2,044 | 10 | 0.325 | 13.0 | 0.436 | 20 | 0.559 | 23.5 | 0.237 |
|  | 1994 | 5,455 | 29 | 0.356 | 45.9 | 0.171 | 50 | 0.690 | 63.3 | 0.108 |
|  | 1995 | 2,075 | 13 | 0.450 | 16.2 | 0.330 | 28 | 1.035 | 38.4 | 0.178 |
|  | 1996 | 1,856 | 8 | 0.520 | 11.0 | 0.472 | 17 | 0.617 | 21.9 | 0.270 |
|  | 1997 | 2,413 | 12 | 0.375 | 16.2 | 0.402 | 20 | 0.625 | 26.7 | 0.228 |
|  | 1998 | 5,334 | 33 | 0.448 | 47.8 | 0.129 | 65 | 0.801 | 89.9 | 0.072 |
|  | 1999 | 1,823 | 9 | 0.389 | 11.9 | 0.493 | 18 | 0.897 | 23.5 | 0.250 |
|  | 2000 | 766 | 4 | 0.359 | 6.1 | 0.720 | 17 | 2.225 | 20.2 | 0.304 |
|  | 2001 | 2,274 | 11 | 0.487 | 16.6 | 0.437 | 26 | 0.492 | 37.7 | 0.189 |
|  | 2002 | 821 | 5 | 0.522 | 5.7 | 0.834 | 16 | 1.101 | 23.0 | 0.352 |
|  | 2003 | 1,422 | 6 | 0.471 | 9.5 | 0.590 | 32 | 1.412 | 47.5 | 0.193 |
|  | 2004 | 911 | 6 | 0.599 | 7.2 | 0.610 | 13 | 2.088 | 17.0 | 0.368 |
|  | 2005 | 2,543 | 14 | 0.998 | 21.4 | 0.407 | 32 | 1.150 | 42.0 | 0.178 |
|  | 2006 | 734 | 4 | 0.891 | 6.3 | 0.770 | 13 | 1.211 | 15.8 | 0.395 |
|  | 2007 | 3,290 | 30 | 0.675 | 55.4 | 0.102 | 55 | 0.632 | 93.6 | 0.066 |
|  | 2008 | 843 | 8 | 1.032 | 12.8 | 0.629 | 38 | 4.145 | 50.7 | 0.191 |
|  | 2009 | 2,448 | 19 | 0.667 | 26.7 | 0.225 | 57 | 2.405 | 81.9 | 0.092 |
|  | 2010 | 6,820 | 71 | 0.583 | 132.4 | 0.055 | 121 | 0.854 | 198.7 | 0.041 |
| Influenza | 2014 | 726,312 | 1052 | 0.085 | 1565.9 | 0.188 | 958 | 0.091 | 1482.0 | 0.004 |
|  | 2015 | 679,850 | 685 | 0.086 | 890.3 | 0.203 | 624 | 0.069 | 848.0 | 0.019 |
|  | 2016 | 704,020 | 696 | 0.072 | 861.8 | 0.224 | 376 | 0.043 | 480.1 | 0.063 |
|  | 2017 | 632,353 | 551 | 0.046 | 712.9 | 0.258 | 659 | 0.047 | 934.2 | 0.008 |

Both approaches had their lowest accuracy on three high incidence dengue seasons: 1994, 2007, and 2010 (Table 3; Fig. 1). The average scores for these years range between 0.041 and 0.17 across the NobBS and benchmark approaches, falling clearly below the rest of the years in performance. These scores not only reflect unusually poor point estimate predictions as judged by rRMSE, but also the finding that the predictive distribution for weeks in these years for both approaches rarely included the true value of interest (a consequence of dramatic over- or underestimates).

### Moving window sizes

We initially used moving windows of 104 weeks for dengue (a longer time series) and 27 weeks for ILI (a shorter time series) to leverage a large number of historical training weeks to train nowcast estimates. Moving windows allow for stable estimation of the recent delay distribution as information from very old and potentially less relevant weeks are forgotten. The size of the moving window reflects how quickly and smoothly changes in the data should be realized by the model: longer moving windows tend to produce smoother estimates, but the model may be less sensitive to abrupt changes in the data (e.g. changes in how quickly cases are reported during an outbreak) or shorter-interval secular trends, e.g. seasonality.

While we chose long moving windows to capitalize on data availability, these considerations may affect the choice of moving window size and nowcast performance, depending on the data. In light of this, we experimented with moving windows of different lengths to assess the impact on nowcast performance with dengue data. We tested moving windows of 5, 12, and 27 weeks (approx. 6 months). A 5-week moving window produced substantially lower accuracy nowcasts (rRMSE = 7.381) with several steep case overestimates in 2007-08 and 2010 (Fig. S6A). However, accuracy metrics for moving windows of 12 weeks or longer were similar to those using the full 104 week window (range in rRMSE: 0.6-0.655; average score: 0.35-0.37) (Table S4; Fig. S6). While shorter moving windows often produced more accurate estimates of the reporting delay probability (Fig. S7A), the estimated variance of the random walk process varied dramatically from week-to-week (Fig. S7B) resulting in more dramatic overestimates at certain periods of extreme time-varying delays, suggesting a trade-off between delay estimation accuracy and more stable estimates of weekly cases.

## Discussion

We introduce a new approach for Bayesian nowcasting and demonstrate its application in two disease contexts with different reporting systems, outperforming an existing method in terms of point estimate (reduced RMSE) and probabilistic (higher logarithmic score) predictive performance. In particular, NobBS performs well even when the delays in case reporting change over time. We further demonstrate an important trade-off related to moving window sizes for delay distribution estimates; short windows improve the real-time characterization of the delay distribution but are susceptible to over-estimating that variability, potentially decreasing nowcast accuracy. Lacking any disease-specific parameterization, and relying only on historical trends of case reporting as input, this approach can be immediately adapted in a variety of disease settings.

Across diseases, NobBS outperformed the benchmark approach on accuracy of uncertainty estimates, and produced comparable or better point estimates. For the subset of weeks in which both models could produce forecasts (weeks with at least one case initially reported), point estimates for NobBS were substantially more accurate than the benchmark model for dengue cases (rRMSE improved by 300%) and slightly less accurate for ILI cases (rRMSE decreased by 19%). However, analysis of the probability distributions of the nowcasts revealed a much more substantial difference; the average score for NobBS was approximately twice as high for dengue and more than 10 times as high for ILI cases (Table 1). This indicates that the NobBS approach assigned much higher probability to the actual outcome, even though point accuracy was somewhat lower for the ILI cases.

While utilizing a similar modeling structure for case reporting delays as the benchmark model (9), NobBS introduces a simple dependency between case counts over time; that is, changes in case counts between weeks are assumed to be related via a first-order random walk process on the logarithmic scale. This feature is critical in the context of infectious disease transmission, where the number of true infections in a given week mechanistically depends in part on the number of true infections in previous weeks due to the infectious process, whether the pathogen is transmitted directly or by vectors (18). Hence, variations of autoregressive models are common in disease forecasting(19, 20). When reporting delays are time-varying, as is often the case in epidemics(17), we show that the NobBS approach is less accurate compared to its performance in a stable delay distribution, but still shows improvement over the benchmark approach likely because the NobBS approach is informed by the number of cases experienced in previous weeks, not just the delay distribution, making it more robust to larger fluctuations.

The accuracy of predictions is related at least in part to the number of cases reported to the surveillance system in week 0. When a larger proportion of cases were reported with no delay, as was the case for ILI compared to dengue, the point estimate accuracy was higher. This is not surprising, as a large fraction of true cases reported initially leaves fewer cases left to predict.

We observed greater volatility in the nowcasts when the initial number of cases reported increases suddenly from low values. Two weeks in the dengue time series highlight this: August 3, 1998 and August 16, 2010. In those weeks, the number of cases initially increased by 16 and 17, respectively, from the previous week after 10 week with an average absolute change of 2.6 and 1.8 cases, respectively. Because this increase is an outlier in the distribution of reporting delays, in particular for delay *d*=0, the model substantially overestimated the true number of cases before correcting the following week. We observed that shorter moving windows either exacerbated this issue (e.g. in 2010) or produced a similar overestimate (e.g. in 1998) (Fig. S6), which appears to be a consequence of the volatility in estimating the variance of the random walk process, despite more accurate estimation of the reporting delay probability (Fig. S7). While the smooth, autocorrelated relationship fit in the NobBS model helps reduce the effect of week-to-week variability in early reporting, it remains a challenge. Users should keep in mind these trade-offs when seeking to apply NobBS to their data.

While NobBS mitigates the effects of a time-varying delay distribution on case estimation, i.e. that the history of cases is leveraged to anchor case estimates to recent values, it does not explicitly model temporal changes in that delay; in other words, the estimated probability of a case occurring with delay = *d* is assumed to apply to all reporting weeks in the moving window. Shorter moving windows can improve estimation of the delay in the presence of changes, but explicitly estimating changes in that distribution may be explored for additional robustness in the presence of systemic changes in reporting. For example, authors in (21) propose a smooth estimate of the time-varying reporting delay distribution using p-spline smoothing. Specifying a time-specific change has also been proposed (9), but empirical identification of a change point in real-time may be challenging or impossible in the context of nowcasting. The challenge that remains in all described approaches is the ability of the model to pick up on changes in the delay distribution that occur quickly, in other words that may otherwise be smoothed out by splines and long moving windows.

Beyond supporting real-time disease tracking by public health officials, NobBS can complement existing disease forecast efforts by providing more accurate nowcasts to forecasting teams in the place of real-time reporting underestimates. For example, teams participating in the Centers for Disease Control and Prevention Epidemic Prediction Initiative (https://predict.cdc.gov) challenges (e.g. FluSight) use initial surveillance data for forecasting because it is the most up-to-date data available (22). NobBS can help account for later revisions to these data and therefore improve prospective estimates as well.

We present an R package, “NobBS,” intended to provide easy and flexible implementation of this approach to a wide audience of public health officials and researchers. This package is currently being finalized and is installable from https://github.com/sarahhbellum/NobBS, and will be moved to CRAN in final form.

## Materials and Methods

### Surveillance Data

We collected data on approximately 53,000 cases of dengue in Puerto Rico and 2.77 million cases of ILI in the United States over a 21-year (1092 weeks) and 3.75-year (196 weeks) period, respectively. Time-stamped weekly dengue data for laboratory-confirmed cases of dengue in Puerto Rico were collected by the Puerto Rico Department of Health and Centers for Disease Control and Prevention. The times used for the analysis were the time of onset as reported by the reporting clinician and the time of laboratory report completion. ILI data originated from the U.S. Outpatient Influenza-like Illness Surveillance Network (ILINet), which consolidates information from over 2000 outpatient healthcare providers in the United States who report to the CDC on the number of patients with ILI. The times used for the analysis were the week of ILI-related care seeking and the week when those cases were posted online in FluView (https://gis.cdc.gov/grasp/fluview/fluportaldashboard.html) as collected in the DELPHI epidemiological data API (https://github.com/cmu-delphi/delphi-epidata). ILI data with delays of more than 6 months occasionally had irregularities, so we restricted the analyses to delays of up to 6 months.

### Reporting Triangle

Delays in reporting are often structurally decomposed into a (*T* × *D*) dimensional “reporting triangle,” where *T* is the most recent week (“now”) and *D* is the maximum reporting delay, in weeks, observed in the data. The data are right-truncated, since at any given week *t*, delays longer than *T – t* cannot be observed. For example, at week *t=T*, only the cases reported with delay *d*=0 are observable; cases reported with longer delays (i.e. 1- or 2-week delays, *d=1* or *d=2*) will be known in future weeks. In Table S5, we present an example of the reporting triangle using ILI data.

For each week *t*, the goal of nowcasting is to produce estimates for the total number of cases eventually reported, *N*_*t*_, based on an incomplete set of observed cases with delay *d, n*_*t,d*_. Since not every *n*_*t,d*_ is observed for a delay *d*, but will be observed at some unknown time point in the future, *N*_*t*_ = sum(*n*_*t,d*_).

Our approach is motivated by modeling the marginal cell counts of the reporting triangle, *n*_*t,d*_ in an adaptation of the loglinear chain ladder method developed in actuarial literature (13, 14).

### Bayesian Nowcast Model

Let *n*_*t,d*_ be the number of cases reported for week *t* with delay *d*. We assume that the underlying cases occur in a Poisson process such that

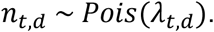

We also allow for extra-Poisson variation, that is, when the variance is larger than the mean and a negative binomial process (of which the Poisson is a special case) is more appropriate. We apply this in the case of the influenza data:

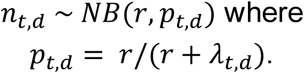

We then model the mean, *λ*_*t,d*_ as a simple log-linear equation

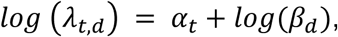

where α_*t*_ represents the true epidemiologic signal for week *t* and *β*_*d*_ as the probability of reporting delay = *d*. In other words, NobBS contains random effects for week *t* and the reporting delay *d*. Exponentiating both sides of the equation,We then model the mean, as a simple log-linear equation 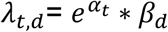.

We place prior distributions on *α*_*t*_ and *β*_*d*_ reflecting properties of each parameter. Since represents a probability vector containing delays = 0, …, *D*, we place on it a Dirichlet prior of length *D*:

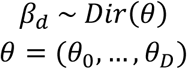

The maximum delay *D* can be identified as the maximum observable delay in the data, which may change as the time series extends, or can be fixed at some value *D* thought to represent a very long delay. In the latter case, *θ*_*D*_ can be modeled as the probability of delay ≥ *D*. For dengue, we choose to fix *D* at 10 weeks, since over 99% of the cases observed in the first two years (prior to producing out-of-sample nowcasts) were reported within 10 weeks. For influenza, we chose *D* to be the longest possible delay within the 27-week moving window, or *D*=26. The implications of choosing a maximum delay *D* within a moving window of *W* weeks means that the nowcast will include all cases arising with delays greater than or equal to *D* but less than or equal to *W*, thus excluding all cases with delays greater than *W* (see the reporting triangle in Table S5).

We place weakly informative priors on θ representing a small number of hypothetical total cases (n=10) distributed across delay bins, loosely representing the probability of reporting delays for each delay *d* observed in the first two years of data for dengue and the first 6 months of data for ILI (training periods). As a sensitivity, we also placed weak priors on θ treating all delays with equal probability, but there was no material difference in the results (Table S6).

We allow a dependency between successive α_*t*_ ‘s to capture the time evolution and autocorrelation of cases from week-to-week, commonly exhibited by epidemic curves. We therefore model α_*t*_ as a first-order random walk:

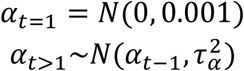

Because *α*_*t*_ is in natural log form, this constitutes a geometric random walk.

We place weakly informative priors on the precisions of the Normal distribution, 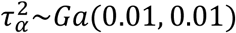. For the negative binomial stopping-time parameter, r, we place an informative Ga(60,20) prior to reflect belief that the process deviates moderately from the Poisson.

Models were compiled in JAGS on R (v 3.3.2) producing 10,000 posterior samples. Trace plots were visually reviewed for convergence.

### Nowcast Estimates

We produced weekly nowcasts beginning with the 27^th^ week (influenza) and 104^th^ week (dengue) and through the final week of the series. This resulted in 989 weekly out-of-sample estimates of dengue cases and 170 weekly out-of-sample estimates of ILI. The time series of key posterior estimates for both diseases are shown in Fig. S11.

We used a two-year moving window to estimate a stable delay distribution within the window. As a sensitivity, and to gauge the minimum amount of historical information required to produce accurate nowcasts, we also applied moving windows of 5, 12, and 27 weeks (approximately 6 months).

We used as a benchmark for comparison the “nowcast” function of the R package “surveillance” by Höhle and colleagues (described in ref. (9)) designed to produce Bayesian nowcasts for epidemics using a hierarchical model for *n*_*t,d* ≤*T-t*_ | *n*_*t,d*_, or the observed cases conditional on the expected total number of cases. We applied the function assuming a time-homogenous delay distribution and recommended parameterization described by the authors in http://staff.math.su.se/hoehle/blog/2016/07/19/nowCast.html, and for comparability, used the same moving window sizes (27 and 104 weeks) to produce nowcasts over the same time periods.

### Model Performance Metrics

The mean absolute error (MAE), root mean square error (RMSE) and relative root mean square error (rRMSE) are defined, respectively, as:

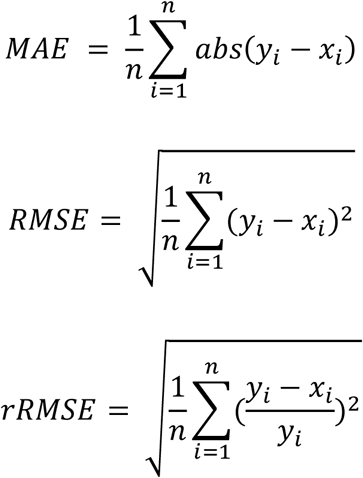

and were used to quantify the accuracy of point estimates, x_i_, compared to true case numbers, y_i_, across the different models at each week *i*.

To quantify the accuracy of the point estimates in capturing the *change* in cases from week *i-1* to week *i*, we computed the mean absolute error of the change (MAEΔ) and the RMSE of the change (RMSEΔ):

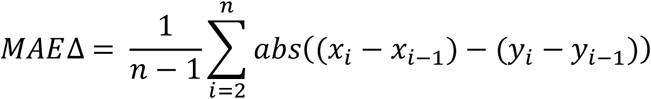

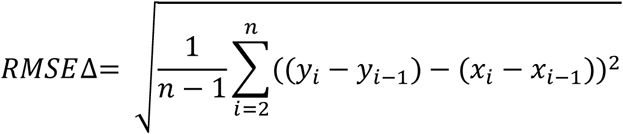

To capture smoothness in predictions from week-to-week, we also calculated the lag-1 autocorrelation of predictions (ρ_a_) and cases (ρ_c_) between week *i* and week *i-1*.

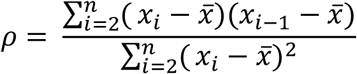

where x = the predicted or true cases at each week *i*.

The logarithmic scoring rule was used to quantify the accuracy of the posterior predictive distribution of the nowcast. Predictive distributions were assigned to a series of bins categorized across possible values of true case counts. We used bin widths of 25 cases for dengue and 1000 cases for influenza, allowing for a larger number of bins for ILI cases based on case ranges of approx. 0-400 for dengue and 4,000-40,000 for ILI. For a predictive distribution with binned probability pi for a given nowcast target, the logarithmic score was calculated as ln(pi). For example, there were 115 cases eventually observed for the week of January 20, 1992. The NobBS nowcast for this week, which assigned a probability of 0.4 to the bin [100,125), thus received a log score of ln(0.4) = −0.92. As in (22, 23), a very low log score of −10 was assigned for weeks in which the predictive distribution did not include the true case value, for weeks in which the bin probability ≤ e^−10^. This rule provides a lower limit (−10) to the score of highly inaccurate predictions.

The average log score across all prediction weeks was computed for all models to assess nowcast performance. The exponentiated average log score yields a nowcast score that can be interpreted as the average probability assigned to the bin corresponding to the true number of cases, and is a metric for model comparison purposes used in several other forecast contexts (22, 23). In this paper, we present the exponentiated average log score and refer to this as the average score.

### Simulated ILI Data

To simulate ILI data with a time-varying probability of reporting delay *d*=0, we drew, for each week, Pr(d=0) from Unif(0.2, 0.9) for all weeks in which the total number of eventually-observed cases exceeded the mean of the ILI series (14,000 cases), and from Unif(0, 0.65) for all weeks in which the total observed case count was less than or equal to 14,000. This probability was used to calculate the simulated number of cases that would be observed with d=0, out of the total number of cases that would be eventually observed for that week. The remaining cases were distributed to other delays ranging from 1-52 weeks using NB(0.9,0.4). This produced a rough approximation for a hypothetical scenario in which cases are reported faster (higher probability of d=0) during weeks with higher disease activity (more cases).

## Acknowledgments

The project described was supported by Grant Number U54GM088558 from the National Institute Of General Medical Sciences and Grant Number 5T32AI007535 “Epidemiology of Infectious Diseases” from the National Research Service Award. The content is solely the responsibility of the authors and does not necessarily represent the official views of the National Institute Of General Medical Sciences, the National Institutes of Health, or the Centers for Disease Control and Prevention.

