## Supplemental Information for "Nowcasting by Bayesian Smoothing: A flexible, generalizable model for real-time epidemic tracking"

### Supporting Information

### Table S1. NobBS Average error for moderate and high absolute changes in initial case reports, compared to the previous week.

| Disease | Threshold | Average error (predicted - actual) for weeks where: Δ initial reports > threshold | Δ initial reports <= threshold |
|---|---|---|---|
| Dengue | Moderate: 5 cases | 61.6 | -1.8 |
| | High: 10 cases | 277.0 | -0.2 |
| ILI | Moderate: 1,000 cases | -92.8 | -38.4 |
| | High: 2,500 cases | -198.9 | -37.7 |

### Table S2. Comparing model performance on influenza reports with constant and non-constant delay distributions.

| Model | | Influenza: Constant delay | | | | | Influenza: Non-constant (time-varying) delay | | | | |
|---|---|---|---|---|---|---|---|---|---|---|---|
| | Period | MAE | rRMSE | RMSE | Average Score | 95% PI coverage | MAE | rRMSE | RMSE | Average Score | 95% PI coverage |
| NobBS | 06/30/2014 - 03/14/2016 | 778 | 0.081 | 1135.2 | 0.172 | 1.00 | 3477 | 0.302 | 4622.7 | 0.06 | 0.93 |
| Benchmark (ref. 9) | 06/30/2014 - 03/14/2016 | 690 | 0.072 | 15559.2 | 0.016 | 0.00 | 7315 | 0.621 | 10300.4 | 8.71E-05 | 0.57 |

### Table S3. Performance measures for estimates of the change in ILI incidence from the previous week, comparing constant and non-constant ILI delay distributions.

| Model | | Influenza: Constant delay | | | | Influenza: Non-constant (time-varying) delay | | | |
|---|---|---|---|---|---|---|---|---|---|
| | Period | MAEΔ | RMSEΔ | $\rho_a$ | RMAΔ | MAEΔ | RMSEΔ | $\rho_a$ | RMAΔ |
| NobBS | 06/30/2014 - 03/14/2016 | 804 | 1268.3 | 0.96 | 1.01 | 4519.573 | 6745 | 0.78 | 3.77 |
| Benchmark (ref. 9) | 06/30/2014 - 03/14/2016 | 758 | 1252.7 | 0.96 | 1.08 | 9437.18 | 14169 | 0.60 | 7.63 |

### Table S4. Select performance measures for dengue fever nowcast model with different moving window sizes.

| Model | Moving window size | rRMSE | Average Score | Correlation |
|---|---|---|---|---|
| NobBS | 5 weeks | 7.381 | 0.368 | 0.275 |
| | 12 | 0.634 | 0.370 | 0.760 |
| | 27 weeks (approx. 6 months) | 0.655 | 0.369 | 0.806 |
| | 104 weeks (approx. 12 years) | 0.600 | 0.349 | 0.84 |

**Table S5. Reporting triangle for June 8, 2015.** The *TxD* reporting triangle decomposes the number of cases, $n_{t,d}$, reported for each week *t* (rows) and each delay *d* (column), up to a delay of *D* = 26 weeks within the 27-week moving window. The goal of nowcasting is to predict the missing (NA) $n_{t,d}$'s.

| | Delay *d* (weeks) | | | | | | | | | | | | | | | | | | | | | | | | | | |
|---|---|---|---|---|---|---|---|---|---|---|---|---|---|---|---|---|---|---|---|---|---|---|---|---|---|---|---|
| **Week *t*** | **0** | **1** | **2** | **3** | **4** | **5** | **6** | **7** | **8** | **9** | **10** | **11** | **12** | **13** | **14** | **15** | **16** | **17** | **18** | **19** | **20** | **21** | **22** | **23** | **24** | **25** | **26** |
| **12/8/2014** | 27487 | 0 | 741 | 504 | 105 | 207 | 40 | 75 | 15 | 24 | 13 | 213 | 6 | 66 | 20 | 6 | 0 | 2 | 0 | 2 | 2 | 0 | 0 | 0 | 0 | 0 | 0 |
| **12/15/2014** | 28964 | 4943 | 1963 | 601 | 232 | 73 | 101 | 94 | 32 | 105 | 372 | 11 | 18 | 18 | 5 | 0 | 0 | 0 | 2 | 25 | 0 | 0 | 13 | 0 | 0 | 0 | NA |
| **12/22/2014** | 32022 | 5648 | 1545 | 298 | 51 | 90 | 31 | 22 | 138 | 398 | 12 | 6 | 13 | 9 | 15 | 0 | 0 | 5 | 23 | 0 | 0 | 8 | 0 | 0 | 0 | NA | NA |
| **12/29/2014** | 32726 | 2372 | 931 | 75 | 236 | 170 | 19 | 141 | 616 | 0 | 5 | 15 | 5 | 4 | 9 | 0 | 2 | 0 | 0 | 1 | 7 | 0 | 0 | 0 | NA | NA | NA |
| **1/5/2015** | 27209 | 2608 | 718 | 330 | 151 | 64 | 184 | 105 | 72 | 42 | 16 | 60 | 11 | 11 | 0 | 11 | 0 | 6 | 4 | 0 | 0 | 7 | 1 | NA | NA | NA | NA |
| **1/12/2015** | 25789 | 3119 | 907 | 390 | 115 | 127 | 454 | 56 | 25 | 22 | 119 | 12 | 12 | 22 | 10 | 28 | 8 | 2 | 0 | 0 | 9 | 0 | NA | NA | NA | NA | NA |
| **1/19/2015** | 28531 | 2502 | 726 | 315 | 250 | 183 | 472 | 24 | 65 | 81 | 15 | 19 | 14 | 8 | 8 | 8 | 0 | 0 | 0 | 6 | 0 | NA | NA | NA | NA | NA | NA |
| **1/26/2015** | 25586 | 3157 | 519 | 263 | 256 | 499 | 54 | 111 | 121 | 40 | 15 | 7 | 8 | 1 | 9 | 1 | 1 | 18 | 14 | 2 | NA | NA | NA | NA | NA | NA | NA |
| **2/2/2015** | 23610 | 2348 | 718 | 465 | 710 | 62 | 145 | 90 | 55 | 25 | 1 | 13 | 0 | 5 | 0 | 0 | 3 | 48 | 0 | NA | NA | NA | NA | NA | NA | NA | NA |
| **2/9/2015** | 20391 | 2808 | 454 | 384 | 490 | 429 | 88 | 51 | 43 | 27 | 41 | 18 | 3 | 3 | 0 | 0 | 18 | 0 | NA | NA | NA | NA | NA | NA | NA | NA | NA |
| **2/16/2015** | 18139 | 1442 | 690 | 451 | 194 | 128 | 71 | 0 | 60 | 29 | 0 | 21 | 2 | 0 | 3 | 21 | 0 | NA | NA | NA | NA | NA | NA | NA | NA | NA | NA |
| **2/23/2015** | 15911 | 1955 | 504 | 306 | 340 | 104 | 7 | 4 | 78 | 31 | 15 | 8 | 0 | 23 | 3 | 0 | NA | NA | NA | NA | NA | NA | NA | NA | NA | NA | NA |
| **3/2/2015** | 15294 | 2037 | 345 | 528 | 103 | 18 | 16 | 163 | 1 | 7 | 5 | 1 | 14 | 11 | 3 | NA | NA | NA | NA | NA | NA | NA | NA | NA | NA | NA | NA |
| **3/9/2015** | 14863 | 1574 | 659 | 151 | 361 | 43 | 24 | 91 | 11 | 3 | 2 | 11 | 18 | 1 | NA | NA | NA | NA | NA | NA | NA | NA | NA | NA | NA | NA | NA |
| **3/16/2015** | 13708 | 2110 | 171 | 441 | 90 | 56 | 111 | 47 | 0 | 1 | 2 | 41 | 43 | NA | NA | NA | NA | NA | NA | NA | NA | NA | NA | NA | NA | NA | NA |
| **3/23/2015** | 13772 | 1372 | 606 | 76 | 186 | 73 | 115 | 0 | 0 | 0 | 8 | 39 | NA | NA | NA | NA | NA | NA | NA | NA | NA | NA | NA | NA | NA | NA | NA |
| **3/30/2015** | 12147 | 1886 | 275 | 252 | 95 | 159 | 62 | 23 | 2 | 16 | 47 | NA | NA | NA | NA | NA | NA | NA | NA | NA | NA | NA | NA | NA | NA | NA | NA |
| **4/6/2015** | 11688 | 1178 | 387 | 111 | 124 | 57 | 18 | 49 | 68 | 37 | NA | NA | NA | NA | NA | NA | NA | NA | NA | NA | NA | NA | NA | NA | NA | NA | NA |
| **4/13/2015** | 9612 | 1158 | 291 | 159 | 164 | 23 | 43 | 44 | 40 | NA | NA | NA | NA | NA | NA | NA | NA | NA | NA | NA | NA | NA | NA | NA | NA | NA | NA |
| **4/20/2015** | 9092 | 1161 | 208 | 136 | 6 | 31 | 30 | 29 | NA | NA | NA | NA | NA | NA | NA | NA | NA | NA | NA | NA | NA | NA | NA | NA | NA | NA | NA |
| **4/27/2015** | 9060 | 993 | 140 | 82 | 172 | 39 | 36 | NA | NA | NA | NA | NA | NA | NA | NA | NA | NA | NA | NA | NA | NA | NA | NA | NA | NA | NA | NA |
| **5/4/2015** | 8702 | 924 | 226 | 213 | 45 | 35 | NA | NA | NA | NA | NA | NA | NA | NA | NA | NA | NA | NA | NA | NA | NA | NA | NA | NA | NA | NA | NA |
| **5/11/2015** | 7558 | 1182 | 241 | 131 | 37 | NA | NA | NA | NA | NA | NA | NA | NA | NA | NA | NA | NA | NA | NA | NA | NA | NA | NA | NA | NA | NA | NA |
| **5/18/2015** | 7015 | 1311 | 482 | 46 | NA | NA | NA | NA | NA | NA | NA | NA | NA | NA | NA | NA | NA | NA | NA | NA | NA | NA | NA | NA | NA | NA | NA |
| **5/25/2015** | 6934 | 799 | 92 | NA | NA | NA | NA | NA | NA | NA | NA | NA | NA | NA | NA | NA | NA | NA | NA | NA | NA | NA | NA | NA | NA | NA | NA |
| **6/1/2015** | 5802 | 642 | NA | NA | NA | NA | NA | NA | NA | NA | NA | NA | NA | NA | NA | NA | NA | NA | NA | NA | NA | NA | NA | NA | NA | NA | NA |
| **6/8/2015** | 4708 | NA | NA | NA | NA | NA | NA | NA | NA | NA | NA | NA | NA | NA | NA | NA | NA | NA | NA | NA | NA | NA | NA | NA | NA | NA | NA |

**Figure S1. The delay distribution (grey) and cumulative distribution (red), in weeks, over the full time series for (A) dengue fever and (B) influenza-like illness (ILI) cases.**

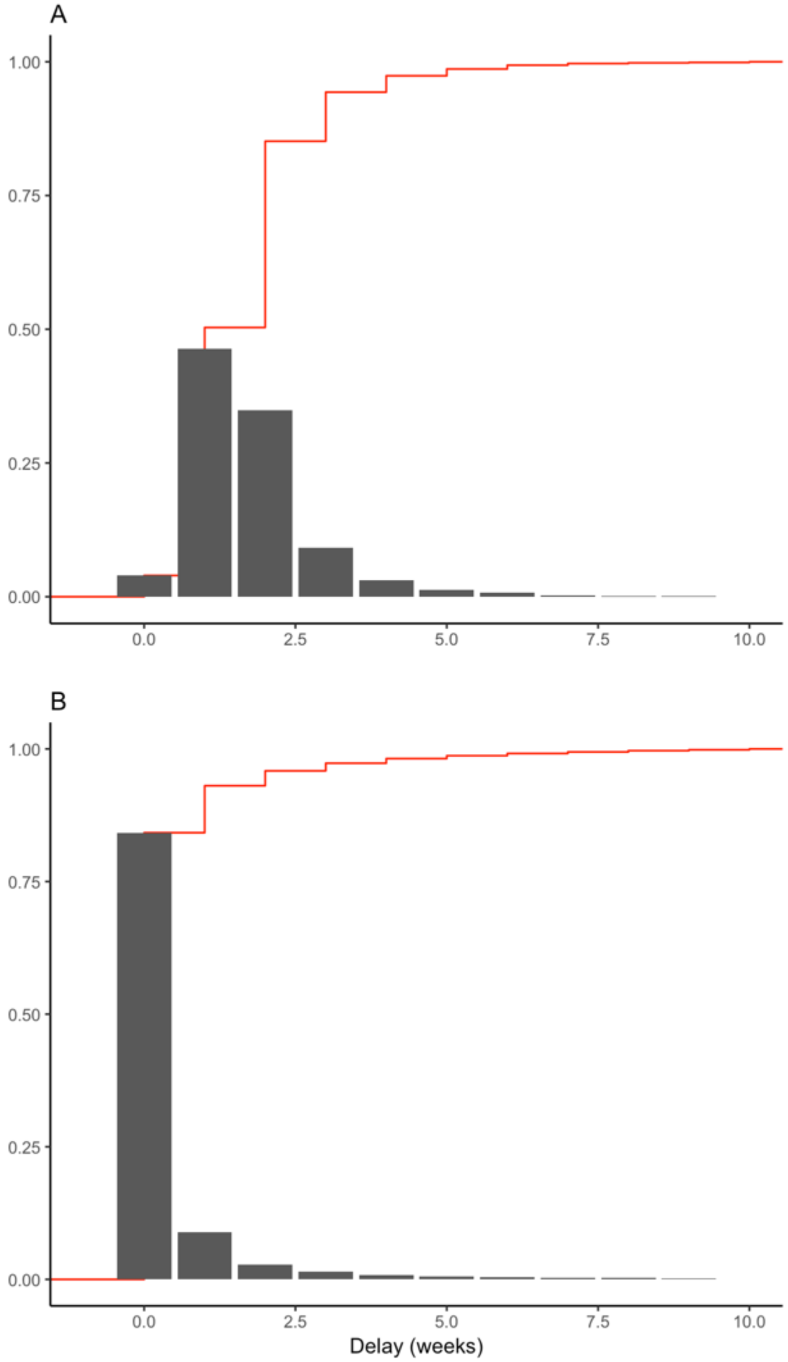

**Figure S2. Comparing (A) the change in initial case reports (from previous week) to (B) the error of NobBS for dengue fever.**

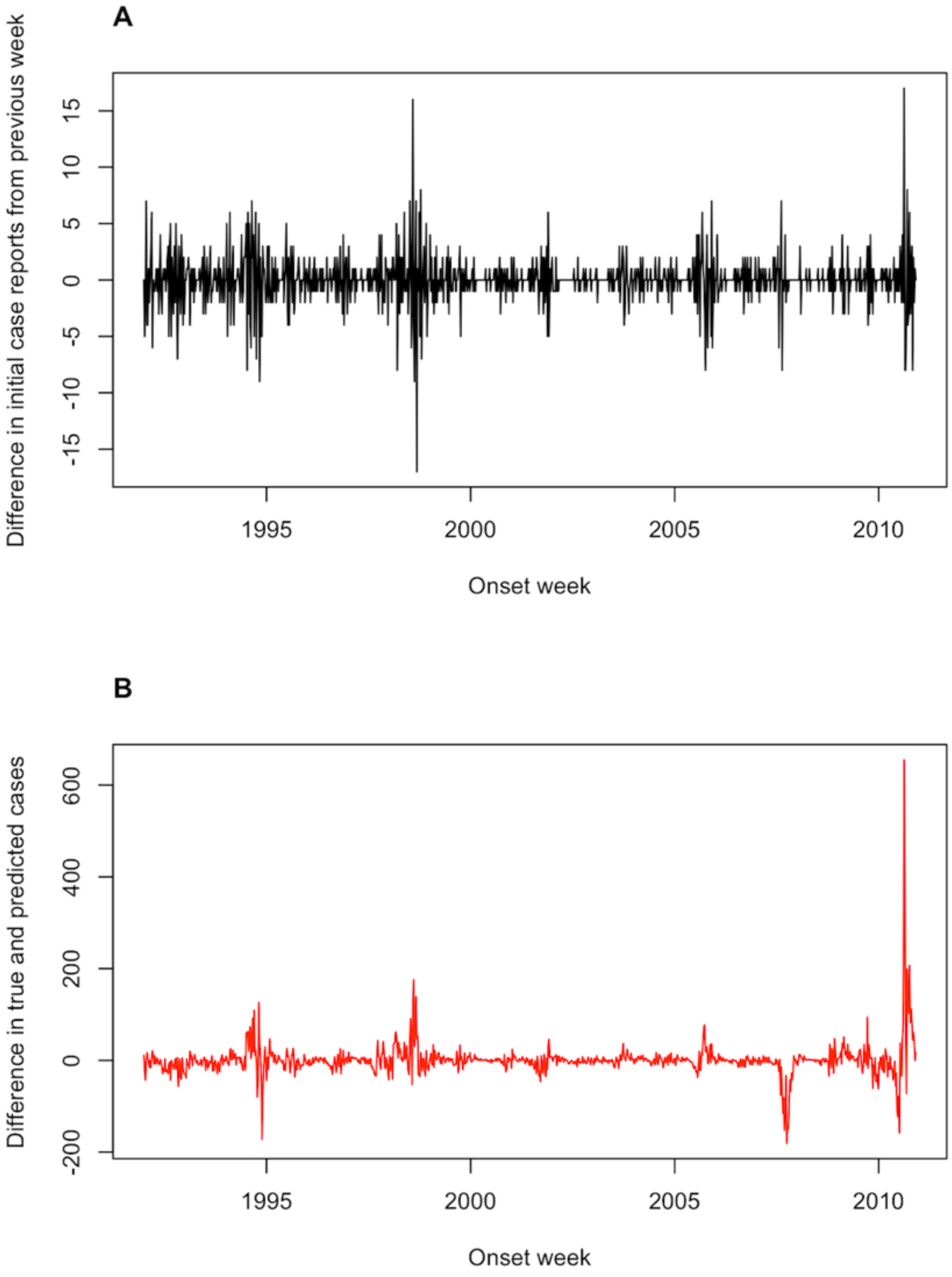

**Figure S3. Weekly reporting delay probabilities for delays up to 17 weeks for (A) dengue fever from 1990-2010 and (B) influenza-like illness from 2014-2017.**

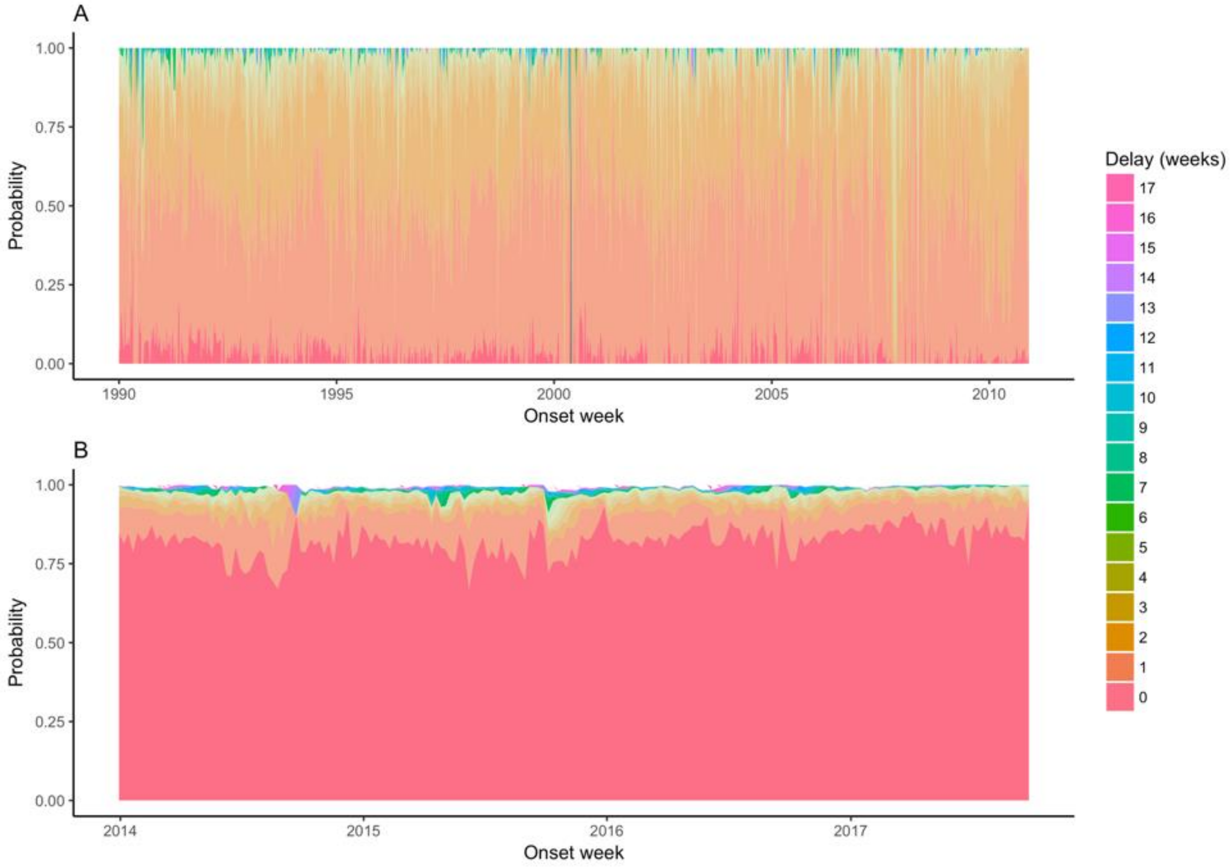

**Figure S4. Weekly ILI nowcasts for June 30, 2014 through March 14, 2016 using a non-constant (time-varying) delay distribution and 2-year moving window.** (A) NobBS nowcasts along with (B) point estimate and uncertainty accuracy, as measured by the log score and the prediction error, are compared to (C) nowcasts by the benchmark approach with (D) corresponding log scores and prediction errors. For nowcasting, the number of newly-reported cases each week (blue line) are the only data available in real-time for that week, and help inform the estimate of the total number of cases that will be eventually reported (red line), shown with 95% prediction intervals (pink bands). For the benchmark approach, the 95% prediction intervals are very narrow and are thus difficult to see. The true number of cases eventually reported (black line) is known only in hindsight and is the nowcast target. The log score (brown line) and the difference between the true and mean estimated number of cases (grey line) are shown as a function of time.

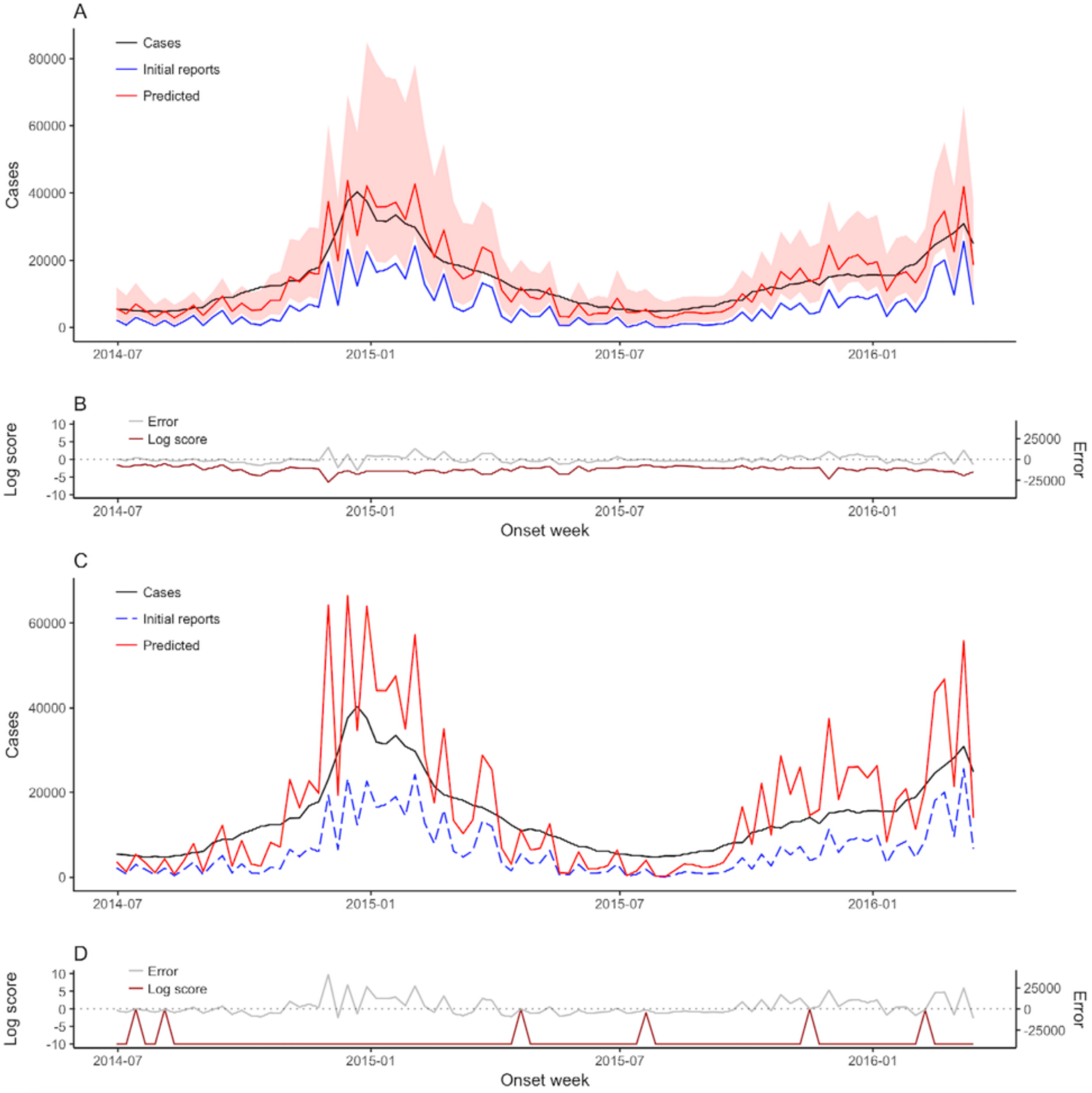

**Figure S5. Comparing the probability assigned to the bin containing the true number of cases (y-axis) to the true number of cases (x-axis), for weekly dengue fever nowcasts using NobBS.** Vertical dashed lines in grey are used to visualize the bin width of 25 cases.

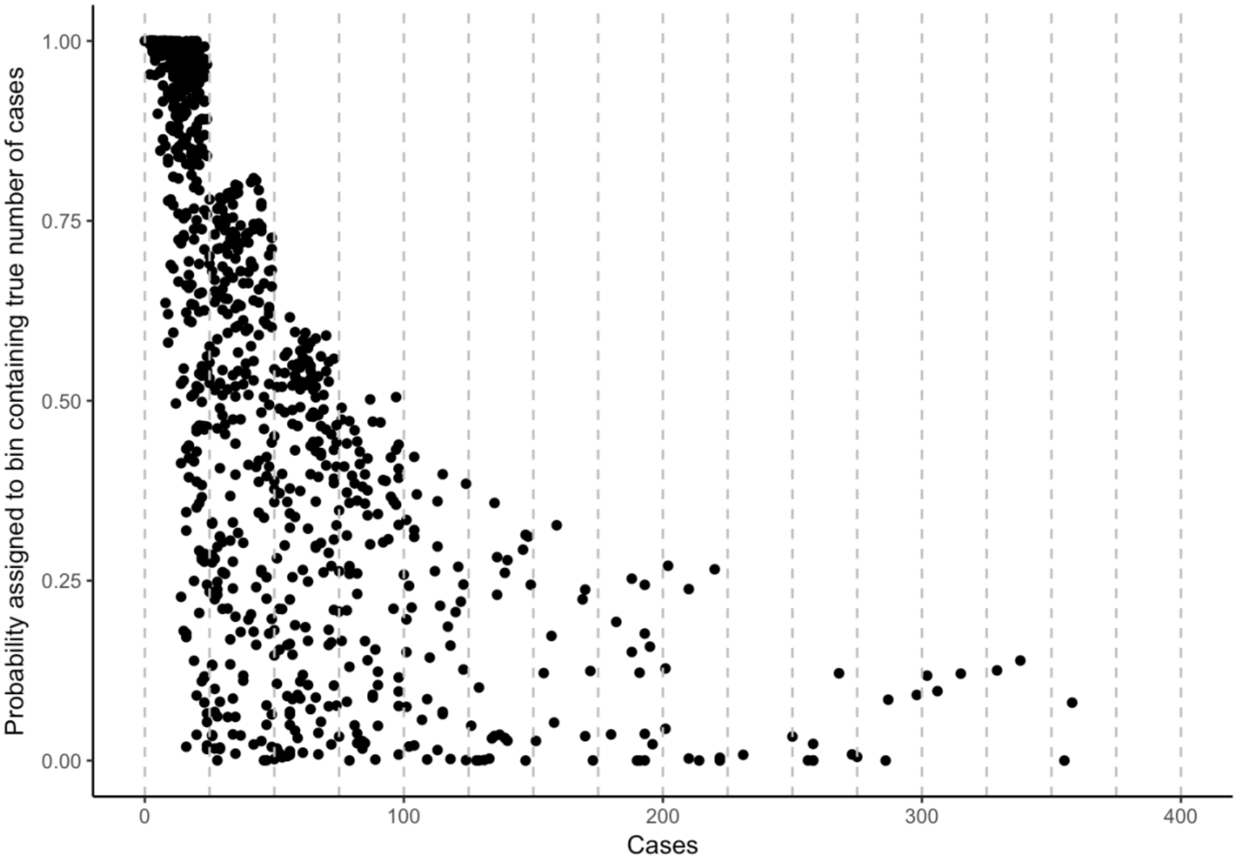

**Figure S6. Weekly NobBS dengue fever nowcasts using (A) 5-week moving window, (B) 12-week moving window, and (C) 27-week (approx. 6 month) moving window.** Plots are zoomed in the y-axis to show the details of prediction.

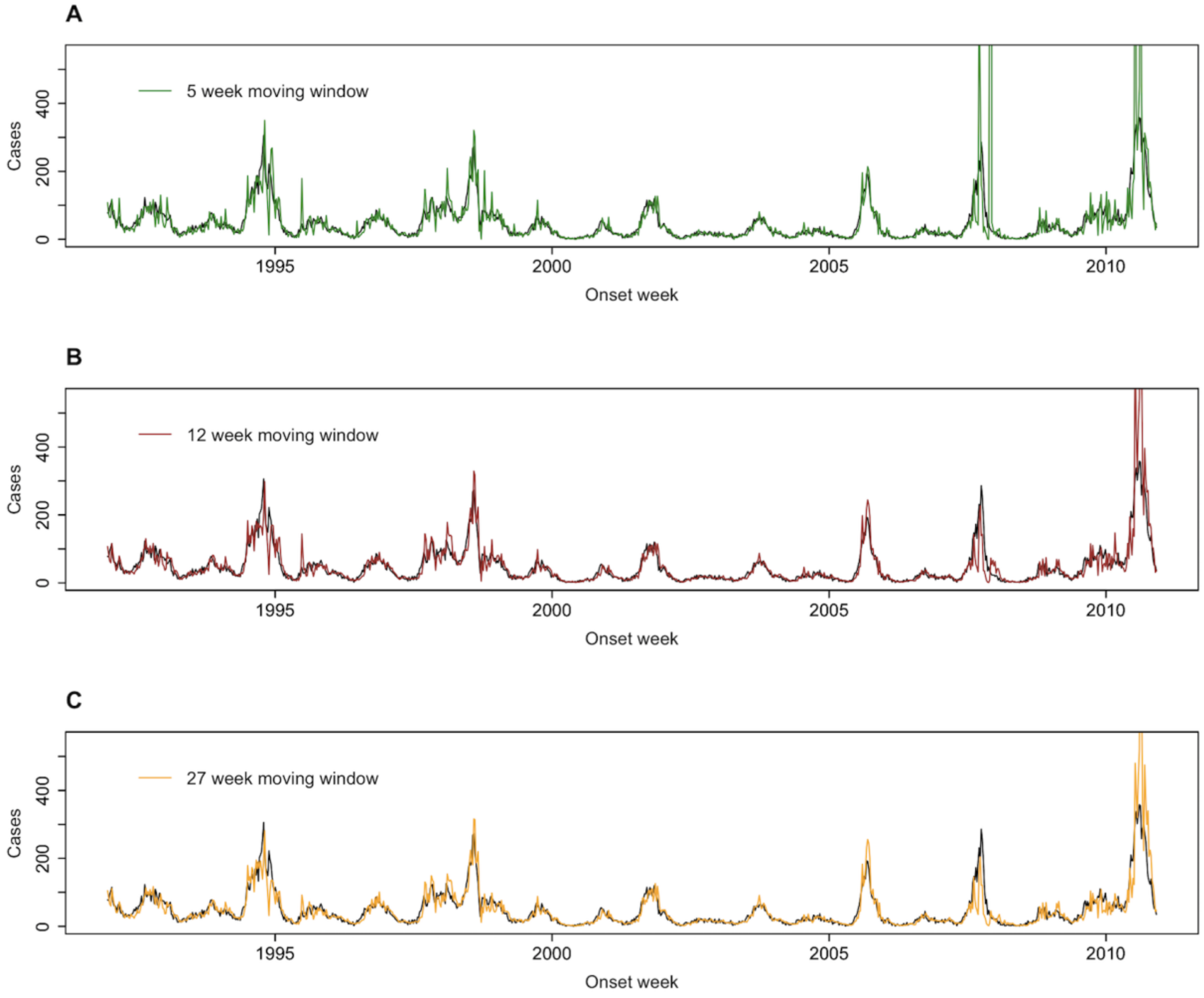

**Figure S7. Comparing the (A) estimated reporting delay probabilities for delay *d*=0 and (B) estimated inverse variance of the random walk, at moving windows of 5 weeks (blue) and 104 weeks (red).** The true reporting delay probability at *d*=0 can be calculated from the data and is shown in grey.

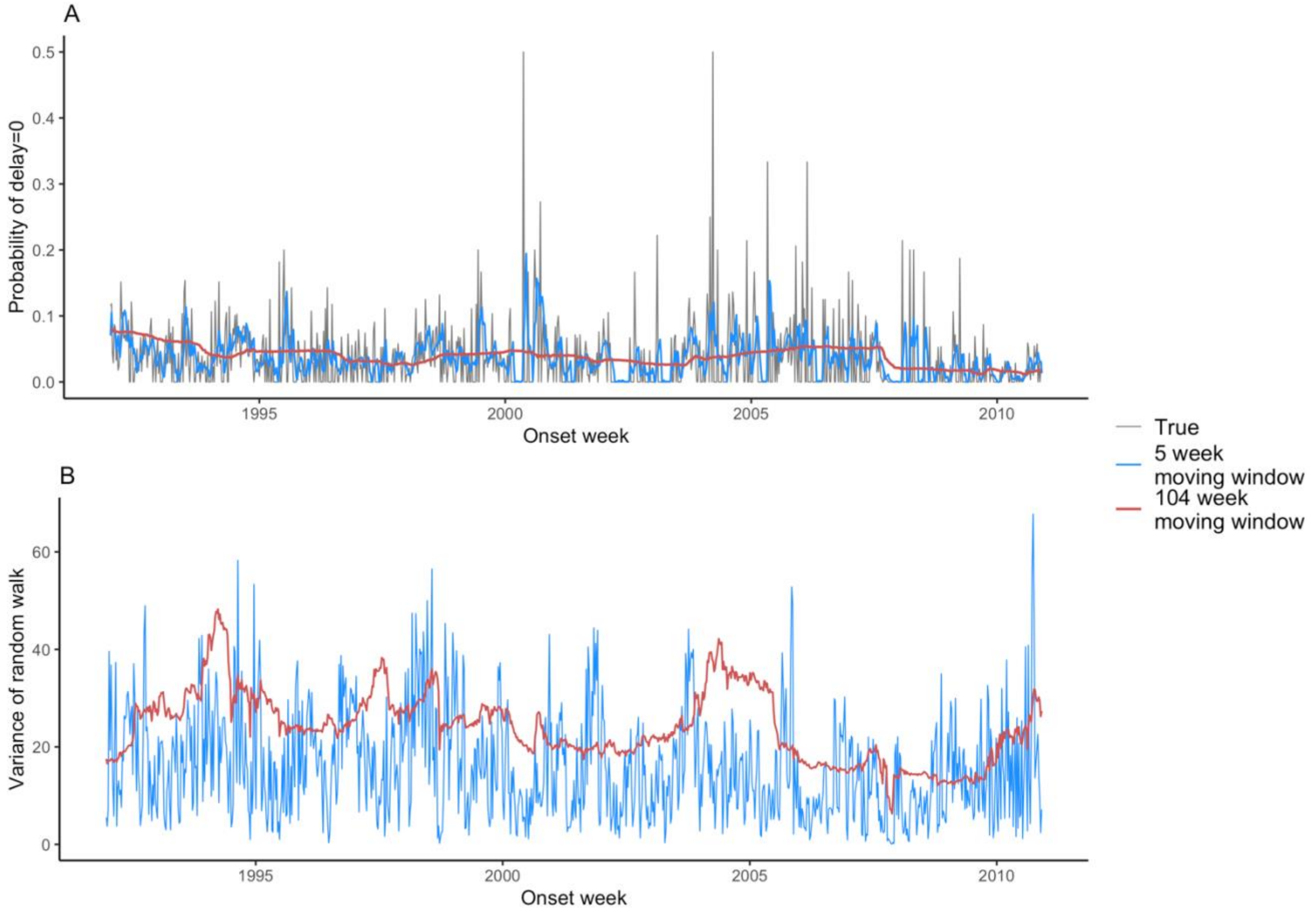
